# Geographic range size is predicted by plant mating system

**DOI:** 10.1101/013417

**Authors:** Dena Grossenbacher, Ryan Briscoe Runquist, Emma E. Goldberg, Yaniv Brandvain

## Abstract

Species’ geographic ranges vary enormously, and even closest relatives may differ in range size by several orders of magnitude. With data from hundreds of species spanning 20 genera in 15 families, we show that plant species that autonomously reproduce via self-pollination consistently have larger geographic ranges than their close relatives that generally require two parents for reproduction. Further analyses strongly implicate autonomous self-fertilization in causing this relationship, as it is not driven by traits such as polyploidy or annual life history whose evolution is sometimes correlated with autonomous self-fertilization. Furthermore, we find that selfers occur at higher maximum latitudes and that disparity in range size between selfers and outcrossers increases with time since their separation. Together, these results show that autonomous reproduction — a critical biological trait that eliminates mate limitation and thus potentially increases the probability of establishment — increases range size.

## Introduction

Why does one species range across an entire continent while its close relative is narrowly distributed (Darwin 1859, p6; Brown et al. 1996; Gaston 2003, ch3)? There are many potential ecological and evolutionary explanations for the enormous variation in species’ geographic distributions. These explanations include species’ geographic location, age, niche breadth, environmental tolerance, competitive ability and key life history traits such as body size, dispersal ability, and mating system (reviewed by Brown et al. 1996; Gaston 2003). Most of these potential explanations have been the subject of intensive research across a wide range of taxa, but none are universally supported. We show that plant species possessing one key trait — the ability to autonomously reproduce via self-pollination — consistently have larger geographic ranges than their close relatives that generally require two parents for reproduction.

To understand the potential impacts of autonomous reproduction, or any trait, on species’ range size, we consider the mechanistic processes underling a species’ range. We envision a species’ range as a set of occupied cells on a grid. Over time, a local population (a cell) might go extinct because individuals cannot tolerate the local environment, because they are outcompeted by another species, or because of demographic stochasticity (Sexton et al. 2009). As some local populations disappear, new populations can emerge by range expansion — the process by which species spread to new geographic locations. To achieve this, individuals must first disperse to new sites. Upon arrival, individuals must tolerate potentially novel biotic and abiotic environmental conditions (Sexton et al. 2009) and, for most sexual species, individuals must find a suitable mate for reproduction and establishment (Baker 1955; Stebbins 1957; Pannell & Barrett 1998). Together, the action of local extinction, recolonization and range expansion generates a species’ range.

Given equivalent amounts of time for range expansion and contraction, consistent differences in geographic range size among species (beyond those expected by stochasticity) are likely due to traits that influence one or more key factors: dispersal, environmental tolerance, and mate availability. For example, increased body size in birds and mammals could increase species’ environmental tolerance by allowing individuals to maintain homeostasis over a wider range of environmental conditions, and body size may increase dispersal ability and home-range size (Gaston 2003, p106-113). Indeed, many studies report that body size correlates positively with species’ range size, although it generally explains only a small fraction of the overall variation in range size (Gaston 2003; Agosta 2013). Similarly, self-fertilization and other forms of autonomous reproduction may affect the basic biological factors that shape range size. Autonomous reproduction directly allows species to overcome limits to range expansion enforced by mate limitation, and may influence range size via indirect effects on dispersal mode (Cheptou & Massol 2009; Hargreaves & Eckert 2014) and environmental tolerance.

There are two primary and opposing hypotheses regarding the effect of selfing, like other forms of autonomous reproduction, on species’ geographic distributions. First, the reproductive assurance provided by selfing is predicted to increase successful colonization and establishment (Baker 1955). Selfers, unlike outcrossing species, are not mate-limited at low population densities and may successfully reproduce when even a single individual lands in a new habitat (Baker 1955; Stebbins 1957; Pannell & Barrett 1998). Under this scenario, species that are already selfing or that evolve selfing will be able to rapidly expand their ranges and occupy habitats that support only small populations or have unpredictable pollinators. Selfers may also be more likely to establish upon colonizing new locations because the capability of autonomous reproduction can shield plants from Allee effects (reviewed in Goodwillie et al. 2005), and because the history of selfing has purged deleterious recessive alleles that would otherwise be exposed in small, isolated populations (Pujol et al. 2009; see Haag & Ebert 2004 for an exploration of this hypothesis in asexual populations). The logical extension of this argument is that selfers will have larger geographic ranges than outcrossers, given similar initial range sizes and amounts of time for range expansion (Henslow 1879, p391; Lowry & Lester 2006; Randle et al. 2009).

An opposing hypothesis is that the limited genetic variation within selfing species may constrain their ability to adapt to many habitats and therefore result in more limited ranges compared to their outcrossing relatives. For instance, self-pollinating populations may have reduced genetic diversity relative to outcrossers (Hamrick & Godt 1996; Crawford et al. 2008). This lack of diversity may limit the niche breadth of selfing populations and prevent them from colonizing and adapting to novel environments (Stebbins 1957; Hamrick & Godt 1996; Crawford and Whitney 2010; Sheth & Angert 2014). Furthermore, although species that reproduce autonomously may initially purge deleterious recessive alleles, they cannot effectively eliminate mildly deleterious mutations, which may accumulate via a ratchet-like process (Heller and Smith 1978; Wright et al. 2013). Competitively superior outcrossing relatives may therefore further diminish the realized niche of selfing species. Ultimately, the evolutionary genetic consequences of self-fertilization may constrain the geographic ranges of selfers relative to outcrossers (Lowry & Lester 2006; Randle et al. 2009).

Here, we test these alternative hypotheses by asking whether, across pairs of sister species of flowering plants from around the world, selfing or outcrossing plants have larger ranges. Data from 194 sister species across 20 genera and generic sections consistently show that selfing species have larger ranges than their outcrossing relatives. Further analyses strongly implicate autonomous fertilization in causing this strong relationship, as it is not driven by traits such as polyploidy or annual life history whose evolution is correlated with the transition to autonomous self-fertilization (Barrett et al. 1996; Barringer 2007, Robertson et al. 2011). Together, our results show that autonomous reproduction — a critical biological trait that influences the probability of establishment — has a major influence on range size.

To identify potential biological drivers of this pattern, we consider two additional questions. First we ask whether species’ latitudinal distributions are explained by mating system. If selfers attain larger ranges by colonizing and establishing at extreme latitudes during interglacial periods (e.g., Jordaens et al. 2000; Griffin and Willi 2014), we expect selfers to be found at higher-latitude locations than their outcrossing relatives. Recently melted glaciers open new habitat for colonization, and the unpredictable pollinator environments at high latitudes may favor establishment of species with autonomous reproduction (Baker 1966; Lloyd 1980). Additionally, the lack of pathogens and competitors at high latitudes relative to tropical regions may permit selfing (for a similar hypothesis in asexual species distributions, see Bell 1982; Glesner & Tilman 1978). We find that selfers do indeed occur at higher maximum latitudes, although we cannot rule out the possibility that a shift in ploidy or life history contributes to this finding.

Second, we ask how differences in range size between selfers and their outcrossing relatives change with the time since their divergence. Continuing range expansion or contraction in selfers will generate an increasing disparity in range size between selfers and outcrossers over time. We find that with increasing divergence time, the range size of selfers increases relative to that of their closest outcrossing relatives, suggesting that range expansion in selfers is ongoing.

## Materials and methods

We identified taxa with a published, species-level phylogeny containing at least one predominantly selfing or functionally selfing species and one predominantly outcrossing species, and with DNA sequence data for at least 50% of the species within the clade available on GenBank (<u>http://www.ncbi.nlm.nih.gov/genbank/</u>) to be used for constructing time-calibrated phylogenies. After removing *Leavenworthia* - a small North American genus in which our phylogenetic model did not converge, we had 20 clades from 15 families whose combined native distributions spanned every continent except Antarctica (see Figure S1 in Supporting Information). On average, clades contained 35 ±7 (±1SE) extant species, 80 ±4.6 percent of which were included in our phylogenies. These time-calibrated, species level phylogenies across a diverse set of plant taxa allow us to test whether mating system influences species’ range size, while controlling for shared evolutionary history.

For the methods and analyses described below, all data and R scripts are available on the Dryad Digital Repository, and software used for each analysis is detailed in Table S1.

### Estimating phylogenies

We reconstructed time-calibrated phylogenies for all 20 clades because most previously published phylogenies were not time calibrated and consisted of only a single topology or consensus tree, making it difficult to incorporate uncertainty into our analysis. Prior to estimating the phylogenies, for each clade separately we downloaded and aligned nrITS sequences for species within the clade from GenBank (Table S1). We simultaneously estimated phylogenetic relationships and absolute divergence times among species in a Bayesian framework (Table S1). Because fossils are not known for our focal clades, we estimated absolute divergence times from the substitution rate for herbaceous and woody plants at the nrITS locus (Kay et al. 2006). The substitution rate was set to a normally distributed prior for herbaceous lineages with mean of 4.13 × 10^−9^ subs/site/yr and standard deviation of 1.81 × 10^−9^, and for woody lineages with mean of 2.15 × 10^−9^ subs/site/yr and standard deviation of 1.85 × 10^−9^.

To accommodate heterogeneity in the molecular evolutionary rate among branches, we used an uncorrelated log-normal relaxed clock model. The prior model on branch lengths was set to a Yule process of speciation. The prior model on substitutions and the number of Markov chain Monte Carlo (MCMC) generations varied by clade (Table S2). Posterior samples of parameter values were summarized and assessed for convergence and mixing (Table S1). After removing *Leavenworthia* (for which the MCMC did not converge, and which we excluded for all analyses) all MCMCs for phylogenies of our 20 clades had minimum estimated sum of squares (ESS) for the posterior >1100, and minimum ESS across all other parameters >600 (Table S2).

For all ensuing analyses, we identified sister species in a subset of 9000 trees from the posterior distribution for each clade. For each sister pair, we recorded the average divergence time and the proportion of trees in which the two species were sister, as a measure of phylogenetic uncertainty. Since our phylogenies sampled, on average, only 80% of extant taxa, these sister pairs may not represent “true” extant sisters, but they are recently diverged species representing independent evolutionary replicates.

### Estimating mating system, ploidy, and life history

We collated 54 studies describing mating systems of species from the clades identified above. Most published studies classified species as predominantly outcrossing, mixed mating, or predominantly selfing. Species were classified as mixed mating when outcrossing rates were between 0.2 and 0.8, or when there was extensive among-population variation in outcrossing rates and traits associated with outcrossing. Exceptions to this classification scheme were species in *Oenothera* sect. *oenothera*, which were classified as either sexual or functionally selfing asexual, due to permanent translocations whereby plants self-fertilize but do not undergo segregation and recombination (Johnson et al. 2009). Sexual *Oenothera* sect. *oenothera* species are partially or wholly self-incompatible, and are assumed to be outcrossing. Different traits are more reliable indicators of mating system in different taxa, and so methods for mating system classification varied among clades, but were generally consistent within clades (described in Table S3). To extend our data set, we occasionally classified taxa that were missing from the primary studies using the same traits and metrics as those used for other species within that clade (Table S3). Only sister pairs with one selfing and one outcrossing species were included in the ensuing analyses, hereafter termed “selfing-outcrossing sister pairs”.

While we focus on mating system, correlated traits such as polyploidy (Stebbins 1950; Barringer 2007, Robertson et al. 2011) and perennial or annual life history (Barrett et al. 1996) may coevolve with mating system. To test whether these traits drive a relationship between mating system and range size, we gathered published information on ploidy and life history when possible. For ploidy, we recorded chromosome counts and classified each species (relative to the base ploidy reported for each genus in the literature) as diploid, polyploid, or mixed when both diploid and polyploid individuals were known. Species’ life histories were classified as annual, perennial, or mixed when both annual and perennial individuals were known. See Table S4 for species’ classifications and sources.

### Estimating geographic range size and latitudinal distributions

We downloaded all known occurrence records for the species in our study from the Global Biodiversity Information Facility (http://www.gbif.org) and filtered for quality by excluding records with coordinate accuracy <100 km, coordinates failing to match the locality description, and taxonomic misidentifications (verified by the authors and taxonomic specialists of each clade). We checked species’ epithets against the most recently published taxonomies and corrected synonyms and spelling errors. To ensure that recent range expansion potentially aided by anthropogenic effects did not influence our results, we included only coordinates from the native range of species. We identified coordinates outside the native species range with published monographs and online databases that report native and invaded ranges (e.g., GRIN database, http://www.ars-grin.gov/).

We used the cleaned occurrence data to estimate species’ range size by dividing the world into a series of rectangular cells by grid lines that follow longitude and latitude (Table S1). We calculated range size as the summed area of occupied grid cells for a given species. To assess whether the ensuing analyses were sensitive to the spatial scale at which species’ ranges are estimated, we calculated range size across a range of cell sizes, 0.05, 0.1, 0.5 and 1 decimal degrees, representing grid cells of approximately 25, 100, 2500, and 10000 km^2^ respectively.

In addition to range size, we quantified three components of species’ latitudinal distributions from the filtered occurrence data: absolute minimum latitude, absolute midpoint latitude (midpoint between minimum and maximum latitude), and absolute maximum latitude. None of the species in our dataset span the equator, so minimum latitude was always greater than zero.

### Analyses

We performed linear mixed effects models to determine whether selfers and outcrossers differ in range size (Table S1). We used natural log-transformed range size as the dependent variable to improve normality of residuals and homogeneity of variance. We treated mating system as a fixed effect and included random intercepts for clades (genus and generic sections) and sister pairs, as well as by-clade random slopes. To incorporate phylogenetic uncertainty into this model, we included a weighting factor for each sister pair equal to the proportion of phylogenetic trees that contained a given sister pair (i.e. the posterior probability of this pair). This weighting allowed us to include all selfing-outcrossing sister pairs, while accounting for the confidence level associated with each sister pair. Additionally, the weighting factor accounts for species occurring in multiple sister pairs across the posterior distribution of trees: the weighting scores of all sister pairs containing a given species never sum to more than 1.

Because the number of sister pairs was highly variable among the 20 clades, we also performed a sign-test (Table S1), with each clade as a single datum. For this test, we took the average sister pair difference in range size (selfer minus outcrosser) for each clade, and asked whether that value was, on average, positive or negative; the null hypothesis was that either was equally likely.

We replicated all analyses across the four range estimates (based on different grid sizes described above) to ensure that our results were robust to the spatial scale at which range size was determined. We also ran all analyses including only selfing-outcrossing sister pairs that did not differ in ploidy (N=127 sister pairs) or life history (N=112 sister pairs) to test whether correlations between mating system and these traits confound our results. We calculated marginal *R*^2^ (proportion of variance explained by the fixed factors alone) and conditional *R*^2^ (proportion of variance explained by both the fixed and random factors) values, and determined significance using likelihood ratio tests with single term deletions (Table S1).

To determine whether selfers occupy different latitudes than their outcrossing sister species, we used the same basic model described above with absolute minimum, midpoint, and maximum latitude treated as response variables in 3 separate models.

To determine whether range size and latitude were affected by divergence time and whether this varied by mating system, we added divergence time and its interaction with mating system as fixed effects to the above model. We allowed clades to have variable slopes by including a by-clade random slope term for divergence time. To meet model assumptions, we log-transformed divergence time.

## Results

The phylogenetic analysis identified 194 sister species that differed in mating system. Within clades, the number of selfer-outcrosser sister pairs ranged from 1 – 68, and their posterior probabilities ranged from <0.01 – 1.0.

Selfing-outcrossing sister species displayed tremendous variation in range size, from having nearly equivalent range areas to differing by > 3 orders of magnitude. Mating system shifts explained a significant proportion of this variation (16-21%) with selfers having, on average, 1.5X – 2X larger ranges than their outcrossing sisters (Table 1; Fig 1). This effect was significant across all spatial scales, and was robust to excluding sister pairs that differed in ploidy and annual/perennial life history (*P≤*0.01 in all cases; see Table S5). Furthermore, a sign test revealed that the predicted clade-average difference in range size (selfing minus outcrossing member of sister pair) was positive (*P*<0.001 in all cases; see Table S6).

**Table 1.** Results of 7 separate linear mixed models analyzing the effect of mating system on species’ range size (estimated at 4 spatial scales) and latitudinal distributions.

| Response | LR | *P* | Marginal | Conditional | <u>Predicted value</u> |  |
| --- | --- | --- | --- | --- | --- | --- |
|  |  |  | *R*<sup>2</sup> | *R*<sup>2</sup> | Selfer | Outcrosser |
| Natural log range size |  |  |  |  |  |  |
| ~25 km<sup>2</sup> grid cell | 12.18 | <0.001 | 0.208 | 0.950 | 1979 | 940 |
| ~100 km<sup>2</sup> grid cell | 11.85 | <0.001 | 0.199 | 0.947 | 6494 | 3224 |
| ~2500 km<sup>2</sup> grid cell | 10.90 | 0.001 | 0.182 | 0.936 | 71,232 | 40,921 |
| ~10,000 km<sup>2</sup> grid cell | 11.15 | <0.001 | 0.159 | 0.930 | 174,312 | 108,597 |
| Absolute latitude |  |  |  |  |  |  |
| minimum | 0.20 | 0.65 | 0.002 | 0.976 | 31.95 | 32.31 |
| midpoint | 1.61 | 0.20 | 0.007 | 0.990 | 37.49 | 36.75 |
| maximum | 3.86 | 0.05 | 0.024 | 0.989 | 43.32 | 41.56 |

Significance of fixed effects was assessed by likelihood ratio tests (LR) using single term deletions. Marginal *R*<sup>2</sup> values are the proportion of variance explained by mating system (fixed effect). Conditional *R*<sup>2</sup> values are the variance explained by mating system and the random effects of clade and sister pair. Predicted values for range size are back-transformed.

**Figure 1.**
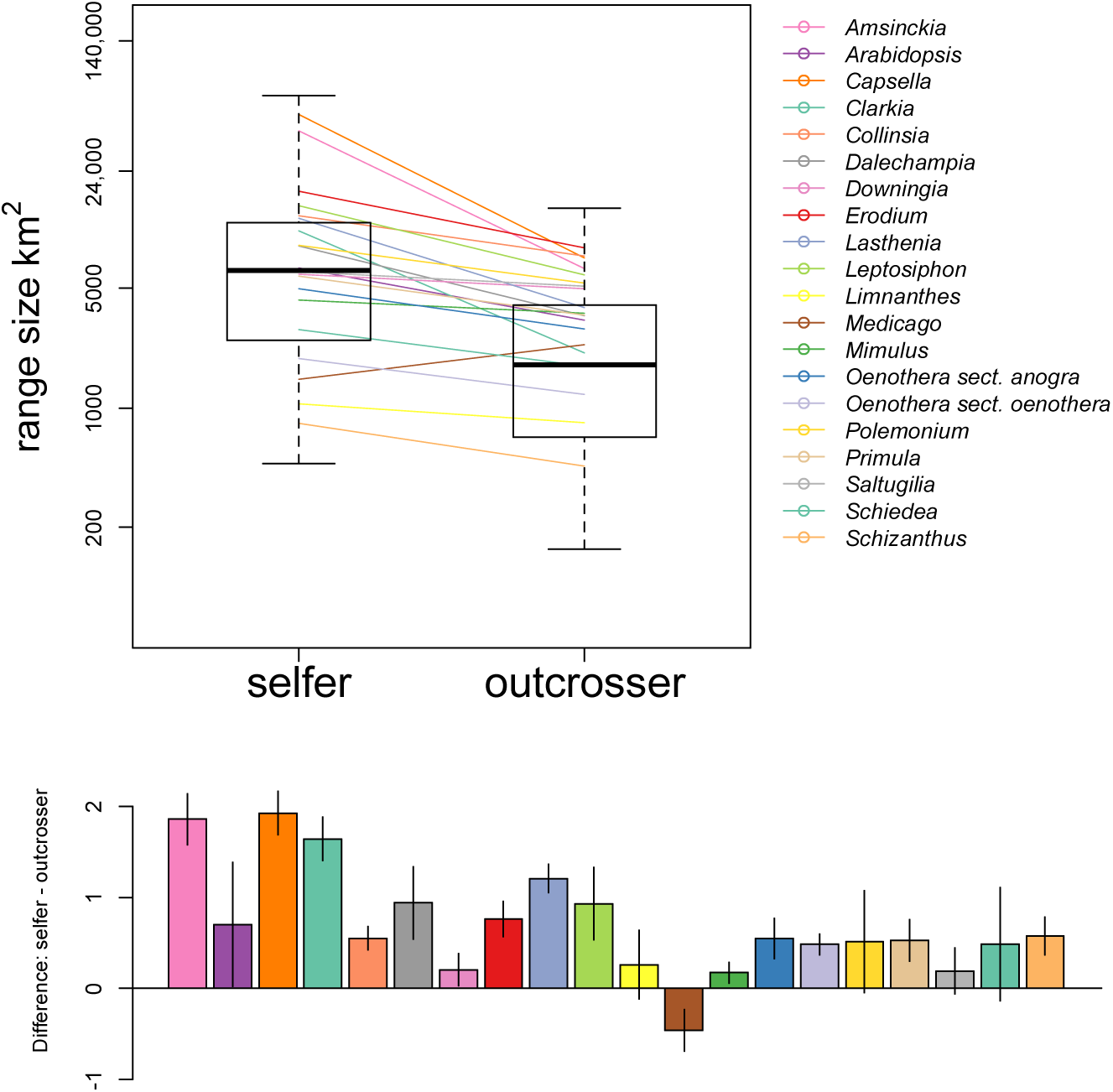
Top panel: Box plots of predicted range size of selfing and outcrossing sister species at the ~100 km^2^ grid cell spatial scale. Colored line segments indicate predicted slopes for each of 20 clades. Vertical axis is natural logarithmic scale (back-transformed km^2^). Bottom bar charts: predicted average sister species difference in range size (ln(selfer)– ln(outcrosser)) for each of 20 clades, with vertical lines representing standard errors. See Table 1 for statistical results.

Mating system also explained the northerly latitudinal distributions of sister species. Selfers had higher maximum latitudes than their outcrossing sister species by about 1 decimal degree or 110 km on average (Table 1; Fig 2A). In contrast, midpoint and minimum latitudes did not vary by mating system (Table 1; Fig. 2B,C). Sign tests revealed that the predicted clade-average difference in maximum and midpoint latitudes (selfing minus outcrossing member of sister pair) was, on average, positive (*P*=0.042 in both cases; Table S6). By contrast, minimum latitude differences did not differ on average from zero (P=0.503; Table S6). When we excluded sister pairs that differed in ploidy or life history, the direction and magnitude of the effect was similar but was no longer significant for maximum latitude (*P*>0.22, see Tables S4 and S5). This loss of significance likely reflects a decreased power, but could also be attributable to polyploidy and annual/perennial life history directly influencing species’ latitudinal distributions.

**Figure 2.**
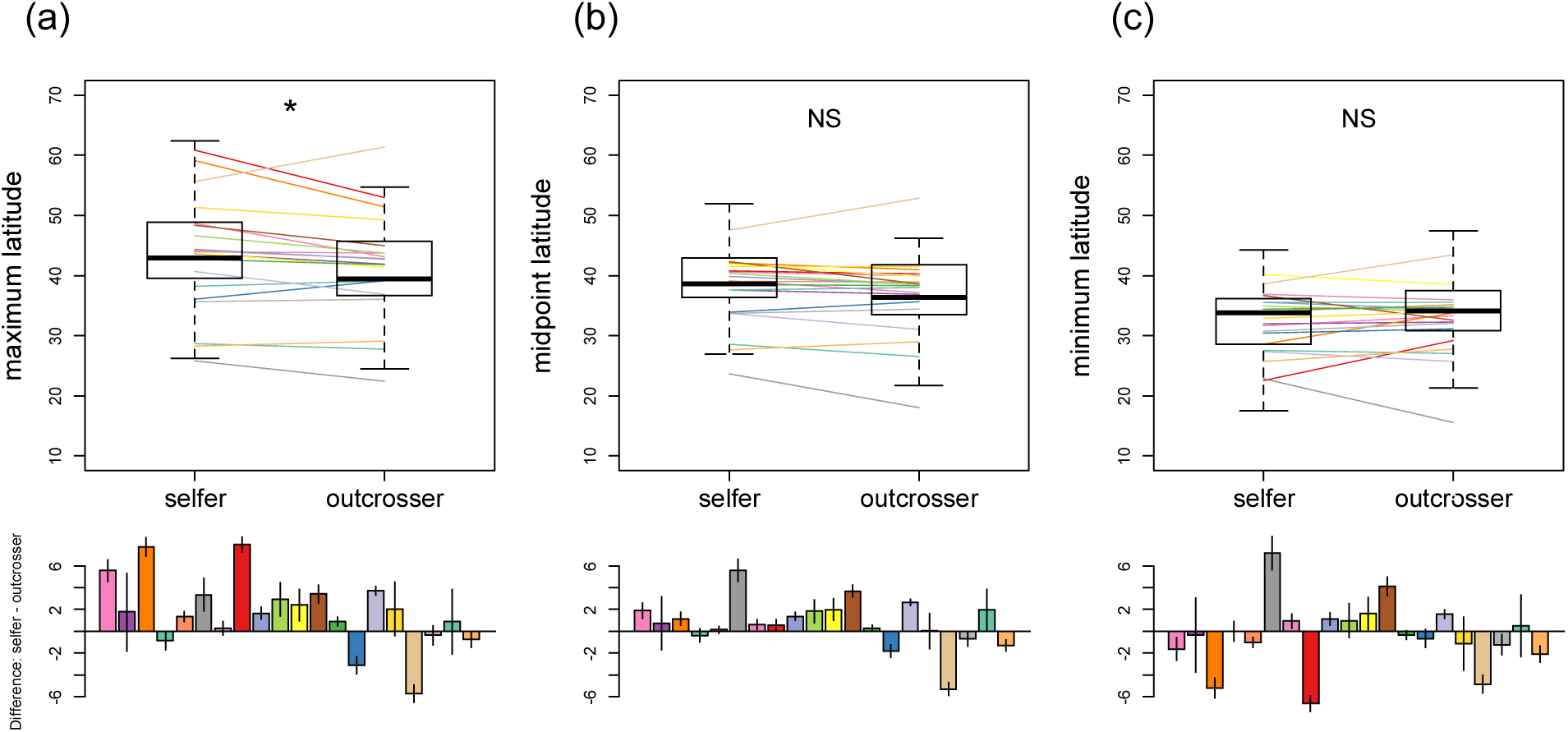
Box plots of predicted absolute latitudinal distributions of selfing and outcrossing sister species, (a) maximum latitude, (b) midpoint latitude, (c) minimum latitude. Colored line segments indicate predicted slopes for each of 20 clades. Bottom bar charts: predicted average sister species difference in latitude (selfer - outcrosser) for each of 20 clades, with vertical lines representing standard errors. * P <0.05. See Table 1 for full statistical results.

With increasing time since divergence, selfing species tended to increase their ranges more than their outcrossing relatives (Table 2; Fig 3). By contrast, neither divergence time, nor the interaction between divergence time and mating system significantly influenced the latitudinal distributions of species (Table 2).

**Table 2.** Results of 4 separate linear mixed models analyzing the effect of divergence time, mating system, and their interaction on species’ range size and latitude.

| Response | Fixed effects |  |  |  |  |  |
| --- | --- | --- | --- | --- | --- | --- |
|  | Divergence time |  | Mating system |  | Time*Mating System |  |
|  | LR | *P* | LR | *P* | LR | *P* |
| Natural log range size | 3.86 | 0.05 | 4.67 | 0.03 | 54.37 | 0.001 |
| Minimum latitude | 2.44 | 0.12 | 0.53 | 0.47 | 2.71 | 0.10 |
| Midpoint latitude | 0.00 | 0.99 | 0.00 | 0.99 | 2.02 | 0.16 |
| Maximum latitude | 0.00 | 0.99 | 3.07 | 0.08 | 2.90 | 0.09 |

Significance of fixed effects was assessed by likelihood ratio tests (LR) using single term deletions. Range size was estimated in ~100 km<sup>2</sup> grid cells; results are consistent across all spatial scales (not presented).

**Figure 3.**
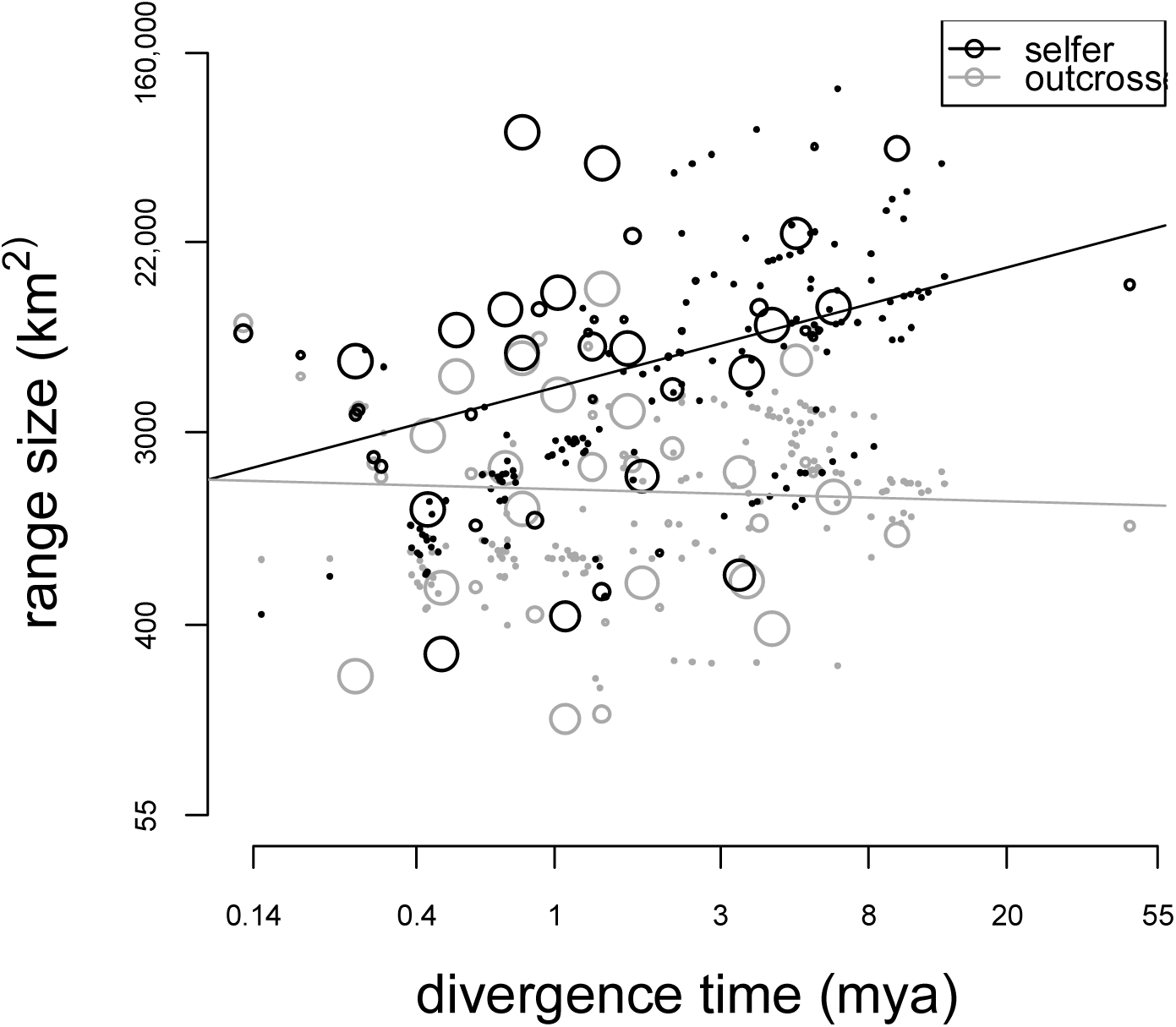
Range size by divergence time for selfing and outcrossing sister species. The size of open circles represents the confidence that species pairs are sisters. The line segments represent the linear regression results for selfers and outcrossers (black and gray lines respectively). Range size and divergence time axes are natural logarithmic scale (back-transformed). See Table 2 for statistical results.

## Discussion

Our analysis across 20 clades (from 15 families) reveals that selfers have geographic ranges on average twice the size of their outcrossing sister species. This difference increases with divergence time between selfing-outcrossing pairs, and may be partially attributable to the colonization of high latitude regions by selfers. Together, our results suggest that the increased colonization ability in selfers associated with the escape from mate limitation (Baker 1955; Stebbins 1957; Pannell & Barrett 1998) allows them to expand and extend their geographic ranges relative to outcrossers (Randle et al. 2009). This effect overwhelms potentially opposing forces that may limit the geographic ranges of selfers.

Henslow (1879) observed a relationship between mating system and geographic range size when he noted that most widespread members of the British flora tended to be selfers. However, it was more than a century until this pattern was systematically tested in four North American plant genera, with mixed results. In *Collinsia*, Randle et al. (2009) found that automatically selfing species had larger ranges than their outcrossing sister species. The same trend was found in *Oenothera* sect. *oenothera*, but it was not statistically significant and relied on different methods (Johnson et al. 2009). In *Clarkia*, the trend was variable, possibly due to poor phylogenetic resolution or to confounding correlated traits such as ploidy (Lowery & Lester 2006). In *Fragaria*, selfing species have larger ranges than outcrossing species (Johnson et al. 2014). The former three genera were included in the present study and supported the overall pattern of greater range size for selfing than outcrossing sister species (Fig 1). It is worth noting that most species examined in past studies have temperate rather than tropical distributions, and 18 of 20 clades in the present study are temperate. Nonetheless, the two clades containing species with largely tropical distributions included here (*Dalechampia* and *Schiedea*) support the pattern of selfers having larger ranges than their outcrossing sisters (Fig 1). As more data become available, it will be valuable to test globally the relationship between mating system and range size.

Our analyses are all correlative, and therefore we are cautious to claim a causative link between mating system and range size. However, we can exclude numerous alternative explanations of this correlation. With additional analyses, we excluded the possibility that life history and polyploidy (traits correlated with selfing) are driving greater range size in selfing species. We also largely rule out two other traits that could drive this correlation: habitat affinity and dispersal ability. If selfing species occupy widely distributed ruderal habitats (Baker 1955; Stebbins 1957) or have greater dispersal abilities than outcrossing species, this could lead to their larger ranges. We account for the former by excluding populations outside the known native ranges of each species, and by restricting the analysis to species pairs with the same life histories (because species that occupy ruderal habitat tend to have annual rather than perennial life histories, Baker 1974). Theory suggests predominant selfers have narrower dispersal distances (Cheptou & Massol 2009; Hargreaves & Eckert 2014), a force going against the results observed here, and therefore different dispersal abilities by mating systems is unlikely to explanation our findings.

We can also largely rule out two potential artifactual explanations for our results. First, because there is more landmass at higher latitudes (particularly in the northern hemisphere), and because selfers occupy higher maximum latitudes, selfers may simply have greater opportunity to achieve larger ranges. However, if this landmass effect drove our observations, it should also apply to the larger-ranged member of outcrossing-outcrossing (O-O) sister pairs. In a separate analysis, we did not find that the larger-ranged member of O-O sister pairs occurs at higher maximum latitudes (see Appendix 1). Second, if sampling effort varied by mating system then range size estimates may be biased. This is not the case – the average number of voucher specimens did not differ by mating system (Appendix 2). Therefore ascribing the larger ranges of selfing species to the escape from mate limitation is consistent with all subsequent analyses, while alternative explanations that we examined are not.

The larger geographic ranges of selfers could, however, reflect greater individual environmental tolerances or species-wide niche breadth. In contrast to the idea that reduced genetic variation in selfers will prevent them from establishing and adapting in new environments (Stebbins 1957; Hamrick & Godt 1996; Crawford & Whitney 2010), we found that selfers had larger geographic ranges, achieved in part by occupation of higher latitudes. Latitude is among the most extreme environmental gradients, with high latitudes experiencing high seasonality and low biotic diversity relative to low latitudes (reviewed in Mittelbach et al. 2007). The ability to persist across a range of latitudes suggests a greater realized species-wide niche breadth for selfers, which may be partially attributable to ploidy and/or life history, or to greater environmental tolerance of selfers versus outcrossers (e.g., Sheth & Angert 2014). Further exploration of the environmental breadth occupied by selfers and outcrossers may uncover axes along which selfers have expanded their realized environmental niches, and generate hypotheses concerning their environmental tolerances.

Like self-fertilization, other forms of uniparental reproduction may allow plants to evade Allee effects during range expansion and achieve larger ranges. Indeed, the ranges of asexual plants are sometimes larger than those of their sexual relatives (Bierzychudek 1985), and asexual species sometimes occur at extreme latitudes (Bell 1982; Bierzychudek 1985). Numerous hypotheses have been put forth to explain the correlation between asexuality and latitude. For example, asexuals may have ‘generalist genotypes’ and broader environmental tolerance (Lynch 1984), or fluctuating biotic interactions with pathogens and competitors in tropical regions may maintain sex in low but not high latitudes (Bell 1982; Glesner & Tilman 1978). Excluding these alternative explanations requires both additional experimental and correlational studies. For example, large ranges in pseudogamous apomicts (plants that, for their own asexual propogation, require pollen which is often not their own (Hörandl 2010)) would argue against our hypothesis that reproductive assurance acts to increase range size.

### The age-range relationship, speciation and extinction

The observation that range size increases with time since most recent divergence has been observed previously, although this effect varies widely across taxa (reviewed in Pigot et al. 2012). We found a similar effect, but only for selfing species (Fig. 3). Although it is tempting to interpret this as evidence of strong directional range size evolution for selfers relative to outcrossers, we caution that the geography of speciation and filtering effects of extinction could also contribute to this pattern. For instance, species that inherit large ranges across speciation events are free to shrink and expand their ranges considerably; in contrast, species that inherit small ranges during speciation will go extinct if their range shrinks substantially. This bias will cause an apparent increase in range size with age, at least for relatively young ages like we consider, because species with decreased ranges were lost to extinction. Thus the geography of speciation and the pace of range size evolution can introduce trends in range size evolution (Pigot et al. 2012). This process could be relevant for speciation events involving selfing-outcrossing pairs (e.g., if selfers commonly arise in small populations via budding speciation); however, these ideas have not been modeled in this context, and therefore the impacts are unclear.

Our finding that selfers have larger ranges than outcrossers, and that ranges of selfing species increase with age, seems at odds with the long-held idea that selfers face high extinction rates relative to outcrossers (reviewed in Igic & Busch 2013). One possible resolution is that selfers do go extinct more frequently, but the ones we observe are those that happen to attain large range size (as in the example above). Another potential resolution is that extinction in selfing plants occurs by rapid extirpations across the entire range, rather than a gradual elimination of populations until the range dwindles and disappears. This could be due to rapid fluctuation of the geographic distribution of selfers, or to stronger autocorrelation of extinction risk across the range. For example, if selfers have less genetic variation (Stebbins 1957; Hamrick & Godt 1996; Crawford & Whitney 2010; Sheth & Angert 2014) and steadily accumulate deleterious mutations (Wright et al. 2013), they may be vulnerable to sudden changes in the biotic or abiotic environments, leading to rapid, range-wide extirpation and extinction. A final potential resolution could be the combination of the phylogenetic scale over which selfing species go extinct (e.g., selfers may give rise to other selfers prior to going extinct) and ascertainment bias on the scale of our study: there are many species, both selfing and outcrossing, whose sister species is not of the opposite mating system and which are consequently not included in our dataset. All of these potential resolutions warrant future research.

### Other contributors to geographic range size

The search for reasons underlying the massive variation in species’ geographic range sizes has a long history that reveals few universal explanations (Brown et al. 1996; Gaston 2003). The results here suggest that mating system is a strong predictor of range size, explaining up to 20% of the variation in species’ geographic ranges. To put this in perspective, body size is among the best-studied correlates of range size and only explains about 6% of the overall variation in the range size of birds and mammals (averaged across studies; Agosta et al. 2013), which is typical of other predictors of range size (Brown et al. 1996). We note that the proportion of variation in range size that we explain is likely an overestimate: we chose clades in which mating system was sufficiently variable so as to potentially explain a reasonable portion of the variation. It is not clear how this ascertainment scheme compares to previous investigations of other traits’ influences on range size.

In addition to mating system, recent shared ancestry may shape species’ geographic ranges (Jablonski 1987; Bohning-Gaese et al. 2006; Martin and Husband 2009). For instance, species generally arise in the same region of the world as their close relatives, which in turn may influence range size, e.g., Rapoport’s Rule (Stevens 1989). Additionally, life history traits that are shared due to recent common ancestry (e.g., dispersal, size, environmental tolerance) may influence species’ ranges (reviewed in Brown et al. 1996; Gaston 2003). In our study, shared ancestry (at the level of genus and sister pair) and mating system together explained 90-95% of the variation in range size.

## Conclusion

Our observation that selfers have larger ranges than outcrossers is consistent with the idea that mate availability at the colonization stage may limit species’ range size. This implies that the ability to find a mate and establish in a novel habitat may have as great an influence on the species’ range as environmental tolerance or interspecific competition. Furthermore, this suggests that traits that increase the odds of finding a mate during colonization (e.g., sperm storage in females), may result in increased geographic range size. Unfortunately, despite the critical role of mate limitation in slowing range expansions (Shaw & Kokko *in press*), studies assessing the impact of mate availability on range size in animals or other taxa are lacking, as much of the last century of research has instead focused on traits influencing dispersal and environmental tolerance (reviewed in Brown et al. 1996; Gaston 2003). We suggest that in many taxa, focusing on traits that encourage mate-finding during colonization may be central to understanding the puzzle of geographic range size.

## Acknowledgements

Boris Igic, April Randle, Sue Kalisz, and the Baker’s Law working group provided stimulating discussions. Anne Worley, Barbara Neuffer, Andress Franzke, Jeremiah Busch, and Justen Whittall provided expert advice on mating systems and phylogeny construction.

## SUPPORTING INFORMATION

Additional Supporting Information may be downloaded via the online version of this article.

